# Can we put Humpty Dumpty back together again? What does protein quantification mean in bottom-up proteomics?

**DOI:** 10.1101/2021.01.25.428175

**Authors:** Deanna L. Plubell, Lukas Käll, Bobbie-Jo Webb-Robertson, Lisa Bramer, Ashley Ives, Neil L. Kelleher, Lloyd M. Smith, Thomas J. Montine, Christine C. Wu, Michael J. MacCoss

## Abstract

Bottom-up proteomics provides peptide measurements and has been invaluable for moving proteomics into large-scale analyses. In bottom-up proteomics, protein parsimony and protein inference derived from these measured peptides are important for determining which protein coding genes are present. However, given the complexity of RNA splicing processes, and how proteins can be modified post-translationally, it is overly simplistic to assume that all peptides that map to a singular protein coding gene will demonstrate the same quantitative response. Accordingly, by assuming all peptides from a protein coding sequence are representative of the same protein we may be missing out on detecting important biological differences. To better account for the complexity of the proteome we need to think of new or better ways of handling peptide data.

## Main

Mass spectrometry-based proteomics has become a key method for characterizing the protein composition of biological samples. The field of proteomics includes a diverse collection of data acquisition and analysis methods, but so-called “bottom-up” proteomics based on proteolysis of proteins into peptide fragments remains the primary strategy for robust surveys of complex protein mixtures. Mass spectra collected from these peptide fragments are then used to infer what proteins were present in the original sample. In the early 2000’s as large-scale peptide identification took off, parsimony was used to assert the set of proteins that could give rise to the peptide data that was observed directly (1,2). As data increased in scale, controlling for false discovery rate (FDR) at the protein level was determined to be a more conservative way to assert protein presence (3).

With the rise in quantitative proteomics, it became desirable to summarize or aggregate peptide quantities into a single value on the protein level. Many strategies have been created to accomplish this, with most assuming that peptides belonging to the same protein will behave similarly. However, based on historical work in protein biochemistry, 2-dimensional gels, and top-down proteomics, it is now estimated that there may be up to 100 proteoforms per protein on average (4,5). This estimate is based on the possible variations that can occur to a protein’s coding sequence or by post-translational modifications (PTMs) to a protein. As most of the amino acid sequence is shared among related proteoforms, a given tryptic peptide can be derived from multiple different proteoforms (**Figure 1**). Once digested in a mixture, the direct connection between a peptide and its originating proteoform(s) is lost, such that the measurements of individual peptides are convolutions of the proteoforms the peptides are present. This issue of conflation is conceptually similar to the problem of haplotype phasing in genomics (5).

**Figure 1.**
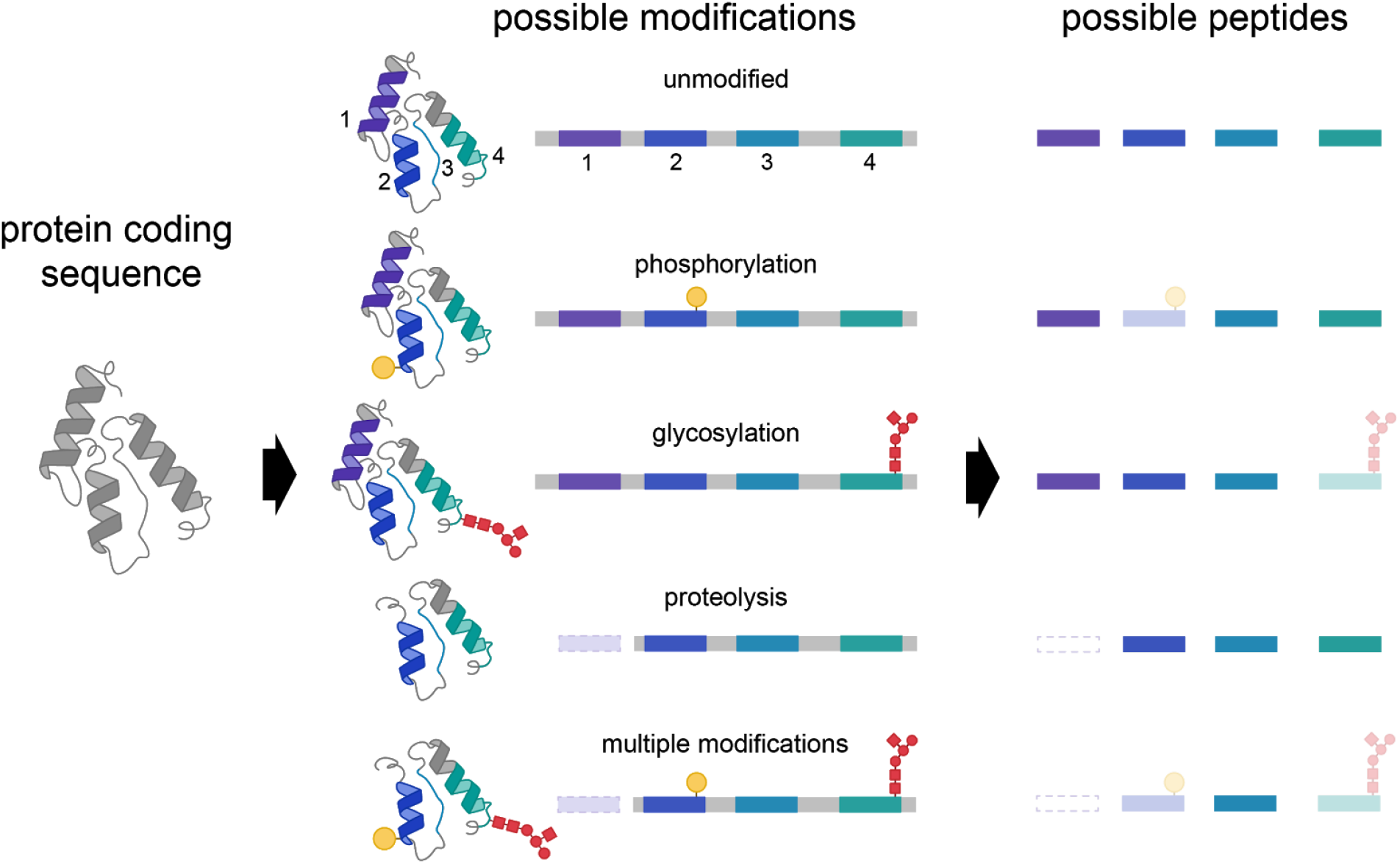
Effect of proteoforms on possible peptide detection. A single protein coding gene can be modified to give rise to dozens or many thousands of proteoforms, including those harboring multiple modifications. After proteolysis, proteoforms yield peptides that may be missed in bottom-up proteomics database searching and data processing.

In quantitative proteomics, our ability to find differences is affected by three parameters: 1) the size of the biological effect, 2) the biological and technical variability, and 3) the number of hypothesis tests that are made within the experiment. Thus, it is important to consider how summarizing peptide quantities at the gene-level will affect these three parameters. The hope is that exploring the conceptual and theoretical effects of summarization to protein-levels will drive improvements in the ability to detect biological effects in a proteomics experiment.

### Rationale for combining peptide measurements to a single protein quantity

The idea of aggregating peptide measurements to the protein level is appealing for interpretation and integration of proteomics data with other data types. Since the beginning of quantitative proteomics, scientists have compared the quantification and coverage of proteomics to the latest gene expression data (6). Intuitively this practice makes sense based on the central-dogma of molecular biology. However, this comparison assumes that for each mRNA transcript there is a single protein quantity for comparison. Despite knowing that there may not be a **single**“protein” derived from the expressed gene, this analysis is standard practice in the field. Such comparisons have demonstrated that the correlation between gene expression and an individual protein measurement is relatively poor (7). While several explanations have been proposed, it is important to note that all experiments were performed with bottom-up proteomics data that has been summarized to a single measurement per protein, even though it is likely that multiple proteoforms exist.

Beyond the proposed ease of biological interpretation, there are technical reasons that make aggregating peptides to a protein level measure attractive. Aggregating peptides mapping to a protein coding sequence into a single measure reduces the number of hypotheses tested, therefore making the analysis theoretically more sensitive to finding protein alterations.

Additionally, by aggregating to a single protein level, whether by averaging or by summing peptide measurements, contributions from outliers or noisy signals are suppressed. This results in the measures having less variation among sets of technical and biological replicates. For example, we can see that in technical replicate runs of cerebrospinal fluid (CSF) digests there is more variability in the peptide level data compared to the protein level values (**Figure 2**). In the case of replicate measures, the reduction in variability is viewed as a positive outcome.

**Figure 2.**
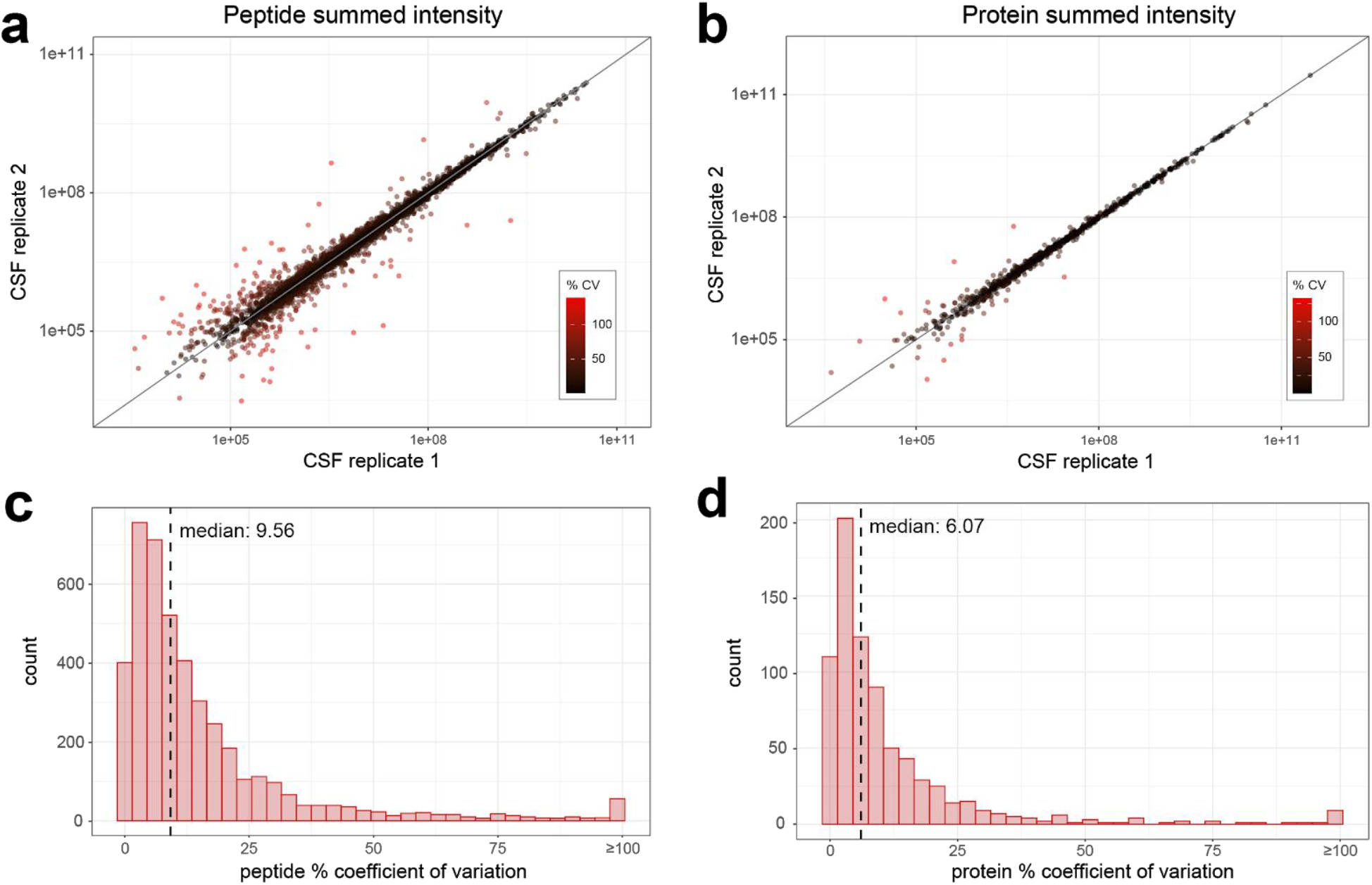
Technical variability is reduced when peptide measurements are combined to a protein measurement. A human cerebrospinal fluid sample digest was analyzed by DIA-MS with 8 m/z staggered windows (4 m/z after demultiplexing). The relationship between a) peptide quantities, or b) summed protein quantities across two replicate instrument runs are plotted, with each peptide colored according to calculated percent coefficient of variation. The distribution of % coefficient of variation for c) peptides and d) summed protein quantities between replicate instrument runs, with the median % coefficient of variation for each indicated by the dashed line.

Another reason to aggregate to a protein level has been to reduce the amount of missing data. With data-dependent acquisition the sampling is stochastic, leading to more missing data at a peptide level if the same precursor is not selected in all the experimental runs. This missing data can have serious implications for the quality of quantitative data. One method to combat this problem is sample multiplexing by isobaric labeling, such as tandem mass tagging peptides. Evaluating large, multiplexed experiments in comparison to label-free approaches, the pattern of the missing data appears to be very distinct, but the macrostructure overall is similar in regard to the relationship between abundance and missing data (8). These multiplexing methods are still limited in the number of samples that can be uniquely tagged, combined, and analyzed at once, and while multiple batches of samples can be acquired, the same peptides are less frequently sampled in different batches compared to proteins (9).

Interestingly, protein groups with greater numbers of peptides observed tend to be statistically different less often than protein groups with fewer peptides (**Figure 3**). Despite the different types of proteomics data, the difference in scale of the data, and using either a sum-based or reference-based quantification, the fold-change consistently trends towards zero. The loss in quantitative significance in proteins with greater coverage is initially counterintuitive. However, because peptide-level quantities are convolutions of proteoform concentrations, a protein with greater peptide coverage will likely span more proteoforms. Unless all those proteoforms change similarly among conditions, aggregating more peptides to a single protein quantity will average away the biological effect. Conversely, if coverage is low, differences in a specific proteoform abundance may not always be reflected with peptide measurements.

**Figure 3.**
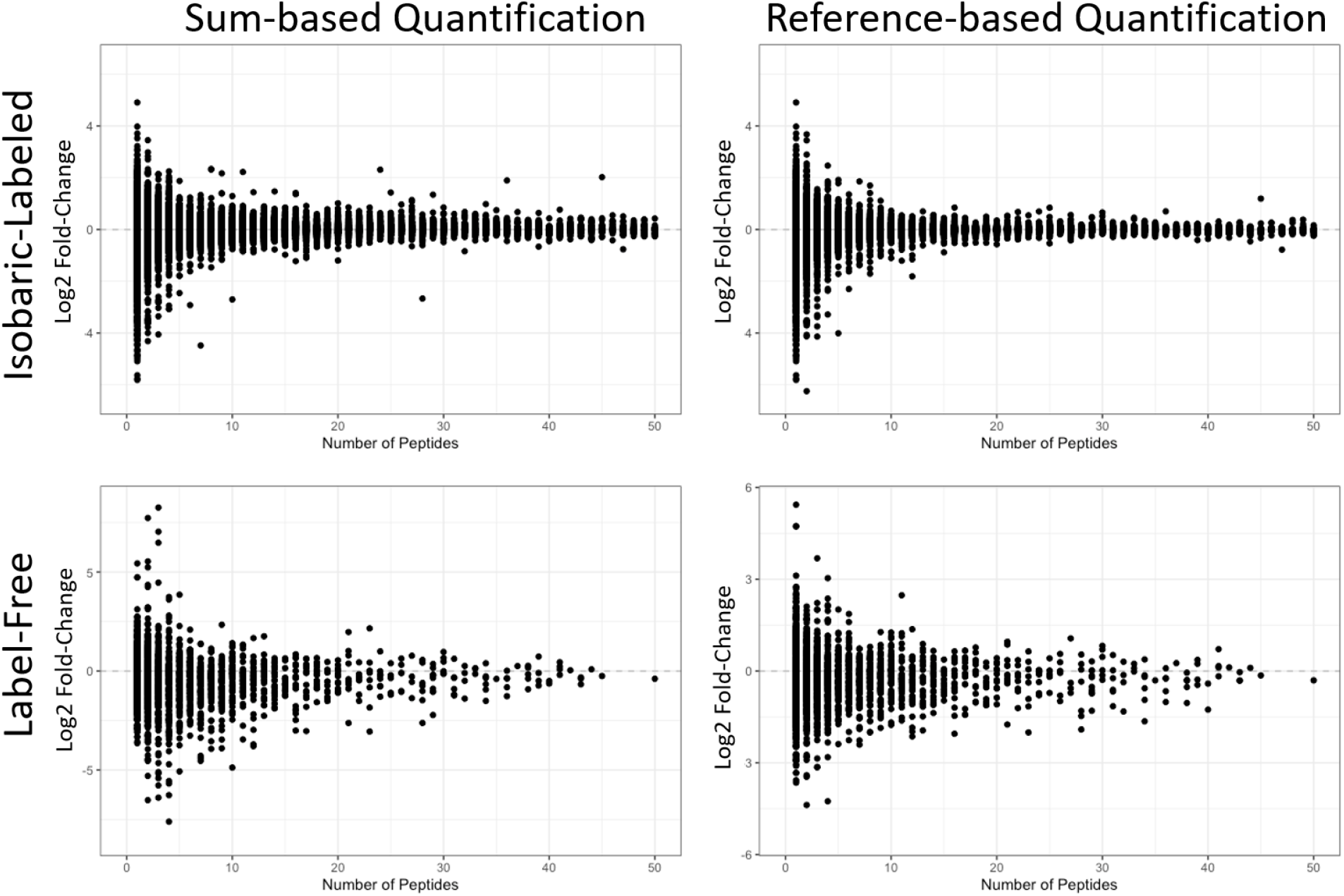
Fold-change estimates for proteins based on the total number of peptides either summed or averaged after reference-based scaling; truncated to 50 for visualization purposes. An isobaric-labeled dataset associated with the Clinical Proteomics Tumor Analysis Consortium (CPTAC) (10), consists of 181,389 peptides mapped to 10,495 unique protein identifiers; proteins ranged from having 1 to 563 peptides associated with them. The log2 fold-change is based on a comparison of tumor residual disease. The second dataset is label free and smaller, based on a Calu-3 cell culture experiment, also publicly available (MSV000079152) (11). This dataset has 15,953 unique protein identifiers, with proteins represented by 1 to 311 peptides. In this dataset the log2 fold-change is based on a Middle East Respiratory Syndrome (MERS) infection to a sham control.

### Limitations of assuming a single quantity per protein coding gene

A consequence of aggregating peptide measures to the protein level is that many key clinical biomarkers of disease do not adhere to the assumption that all peptides mapping to a gene behave similarly. To the extent this is true, statistical power will be reduced in connecting phenotype to proteomic data. For example, amyloid-beta is a peptide derived from the amyloid precursor protein gene. In Alzheimer’s disease a series of cleavage events lead to several shorter soluble forms of amyloid precursor protein (sAPPα, sAPPꞵ), C-terminal fragments (AICD50, CTF 83, CTF 89, CTF 99, p3) and amyloid-beta peptides, which contribute to forming the characteristic plaques observed in the brains of diseased individuals (12). The amyloid-beta peptides can be variable lengths depending on specific cleavage site, but commonly occur as a peptide of either 40 or 42 amino acids (13). In addition to the widely known amyloid-beta 40 and amyloid-beta 42 peptides, over 20 additional amyloid-beta proteoforms have been detected in samples of Alzheimer’s brain samples arising from endogenous cleavage and post-translational modifications (14,15). Knowing that the amyloid precursor protein is heavily processed, it is difficult to determine the origin of many of its tryptic peptides - whether they are derived from an unprocessed amyloid precursor protein, or from one of many processed forms.

If we aggregate all the tryptic peptide measures, we are assuming they are all derived from the unprocessed state, which may not be the most accurate assumption for peptides mapping to amyloid precursor protein. If we look at data from tryptic peptides, we see that some biologically relevant differences would not be accurately represented if our peptide measures are combined to a singular protein level (Figure 4). Specifically, in tryptic peptides mapping to the region of the amyloid beta sequence we observe a different abundance profile compared to tryptic peptides mapping to other regions of the protein. In addition to amyloid beta, phosphorylated tau proteoforms in the cerebrospinal fluid of patients have also gained acceptance as diagnostic biomarkers of disease (16). Additional studies indicate that specific tau phosphosites may be better indicators of disease progression, emphasizing the importance of distinguishing between different pTau isoforms and proteoforms (17).

**Figure 4.**
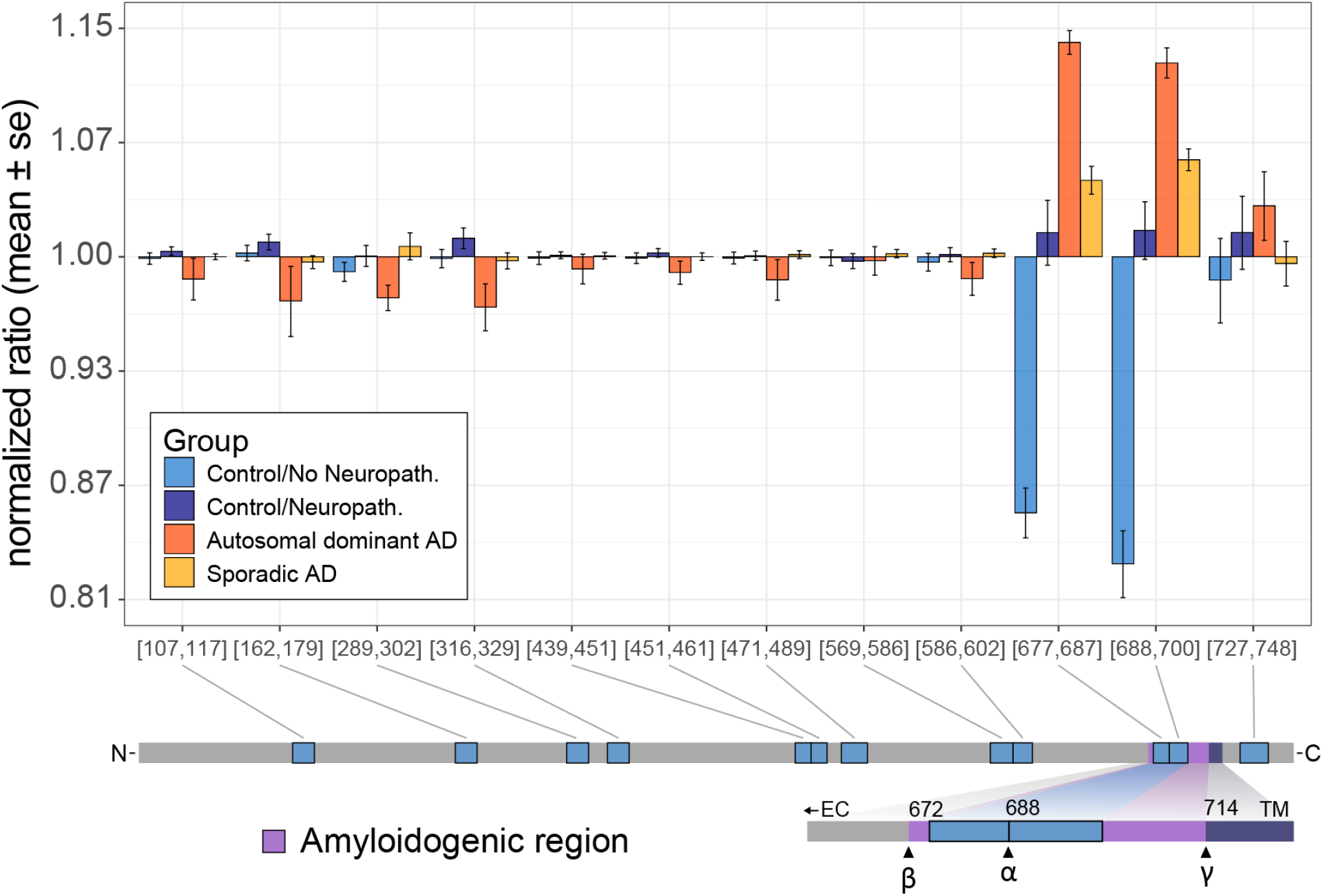
Differential abundance profiles of tryptic peptides mapping to amyloid precursor protein. Three experimental groups of patients were analyzed by DIA-MS; Control/No Neuropath with normal cognitive function and no neuropathologic changes of Alzheimer’s disease including no amyloid accumulation, Control/Neuropath with normal cognitive function and intermediate or severe level of neuropathologic changes of Alzheimer’s disease, Sporadic AD with dementia and intermediate or severe level of neuropathologic changes of Alzheimer’s disease, and Autosomal dominant AD with dementia and intermediate or severe level of neuropathologic changes and an autosomal dominant mutation. For all unique peptides mapping to the amyloid precursor protein sequence, peptide measures are normalized to the mean and the mean & standard error are plotted by group. Based on known protein processing we see that the two peptides with large differences map to the amyloidogenic Aꞵ polypeptide.

Ambiguities due to modified or processed protein biomarkers is not a problem unique to Alzheimer’s disease, but rather is general to human biology and therefore human disease. The products of processing a precursor protein into polypeptides are important markers in diabetes. C-peptide and insulin are both derived from proinsulin, with C-peptide being a valuable measure of insulin secretion and therefore pancreatic beta cell function (18). Proglucagon is processed to form up to nine different polypeptide products, including the better-known glucagon and GLP1. Both polypeptides have distinct roles in metabolism, and both are drug targets for diabetes and obesity (19). Additional examples of this type of processing can be found in the kallikrein-kinin system and coagulation pathways (20). While these examples are well studied, we should not assume that these types of modifications leading to unique biologically relevant proteoforms are uncommon among other less studied proteins.

Along these same lines, a study of cerebrospinal fluid in Parkinson’s disease found that quantification of specific tryptic peptides was differential in affected individuals compared to healthy, age-matched controls. Specifically, peptides in the C-terminal or N-terminal regions of granin family proteins were found to be decreased in Parkinson’s (21). Importantly, the granin family of proteins is known to play a role in regulating secretion and delivery of peptides and neurotransmitters and are known to be processed into a number of derived bioactive peptides (**Figure 5**). As demonstrated in figure 5, if we sum all peptide measures that map to the protein coding sequence of secretogranin 2, then we miss the differences between experimental groups for several of the individual peptides. Instead, aggregating peptides to a single measure per protein coding sequence only accurately reflects the peptide level measurements if all peptides are in agreement (**Figure 5**). In contrast, if we look at peptides detected and quantified from GAPDH protein in the same CSF experiment we observe the same trend across peptides. Interestingly, GAPDH has been observed not to have many proteoforms by top-down analysis (22). Although there are known proteoforms, from the peptides we detect we cannot conclude that only one proteoform of GAPDH is present in our samples. Instead, we can only conclude that all the peptides we detect share the same abundance trend.

**Figure 5.**
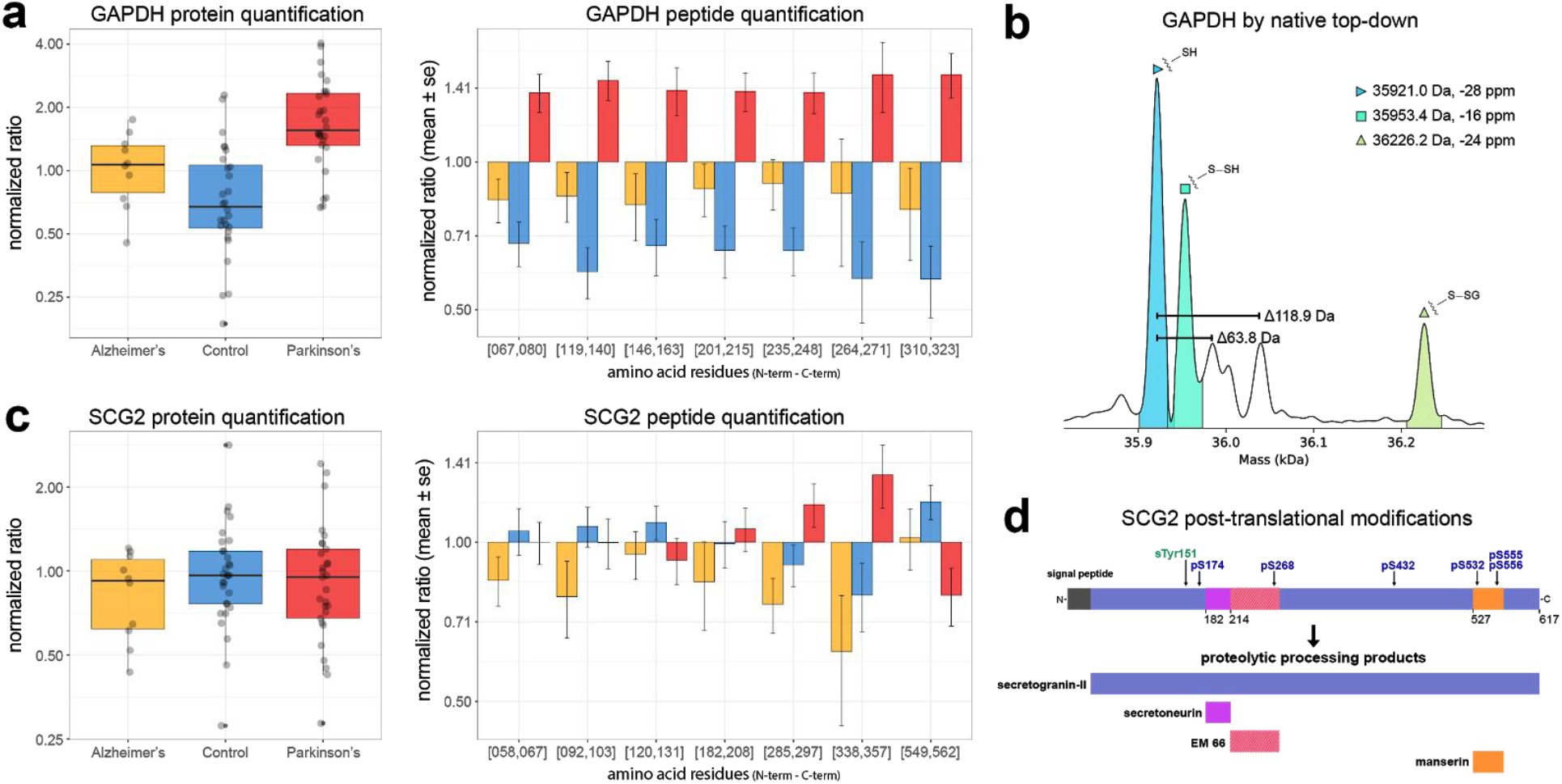
Abundance profiles of tryptic peptides mapping to a) GAPDH and b) SCG2 proteins in cerebrospinal fluid. Three groups of human cerebrospinal fluid samples were analyzed by DIA-MS: Alzheimer’s disease, Parkinson’s disease, and healthy age and sex-matched controls. Unique peptides mapping to the proteins a) GAPDH and c) SCG2 report quantitatively on their relative expression ratios. The protein-level display integrates the mean values from all peptide-level results (box-and-whisker plot at left), with the expression ratio for each individual peptide and the group shown in the bar graphs at right. b) GAPDH has been observed as three proteoforms which form homo-tetramers from human cell lines including HEK-tsa. Intact mass spectra of the monomeric form reveal a canonical form, a persulfide-modified form, and a glutathione-modified form. Reported masses represent average masses and ppm mass error from the calculated theoretical average mass. d) SCG2 is proteolytically processed to produce several peptides, has a sulfotyrosine, and can be phosphorylated at several serine residues.

While bottom-up proteomics is arguably the most common method for characterizing protein mixtures, alternative methods are gaining interest. These include methods that use antibodies and aptamer affinity to recognize a specific protein or protein domain (23–25). Although these methods will usually not have to deal with multiple measures resulting from a protein coding gene, they instead rely on single measurement. It should be noted that any method that constrains complex proteoforms into a single quantitative value per protein coding gene may miss many of the underlying differences.

### Outlook and Future directions

Over the years, this challenge that not all peptides from the same gene or protein group have the same differential abundance has been an important area of research. The approach of several proposed methods focuses on the exclusion of peptide measurements from inclusion in the aggregate protein quantity if they are outliers from other peptide measurements mapping to the same protein coding gene (26–31). While these all demonstrate improved protein concentration estimates, they still only report a single protein quantity and ignore peptides that do not agree with that single value. If those outlier peptide measurements are discarded, then true biological signals may be lost. Another approach taken in previous methods is to try and identify the specific proteoforms present based on peptide quantification across conditions (32–34). While more tolerant to the possibility of having multiple proteoforms present in a sample that lead to altered peptide abundance, the ability to confidently assign peptides to specific proteoforms is all but impossible in bottom-up proteomics.

While the challenge of aggregating peptide measurements may not be solved yet, one thing that is apparent is that we should no longer blindly merge all peptides into a single gene level quantity. A solution to the presence of discordant peptides could be to keep all peptides as independent measurements because it is impossible to accurately merge peptides without detailed knowledge of all proteoforms in the sample. While remaining as true to the acquired data as possible, this strategy may prove to be difficult for interpretation of experiments because the role of individual tryptic peptides in biology may be difficult to infer, especially in less studied systems. Additionally, reduced statistical power for differential abundance testing on tens of thousands of peptides compared to thousands of protein groups will also likely result in fewer significant differences. However, there has been recent work towards integrating top-down proteomics with bottom-up proteomic measurements (35). This strategy could provide higher resolution information about the protein quantity resulting from specific proteoforms present in a sample, which then can be used to determine how peptides could be combined to more accurately reflect those proteoforms present.

An alternative approach could be to combine peptides that both map to the same gene and co-vary across a diverse set of biological groups or conditions, without designating them as specific proteoforms. We need the ability to generate multiple “peptide groups” for each protein group --resulting in 1 to N quantities for each protein where N is the number of peptides. This grouping would require a method that minimized variance and multiple testing while maximizing the biological effect. This approach would not require knowing which proteoforms were present but would still capture quantitative differences observed at the peptide level that would otherwise be eliminated by combining those differences with non-changing peptides within the same gene product. However, this approach could be heavily dependent on having multiple conditions with enough biological replicates and high reproducibility. Additionally, the approach may not be suitable for proteins with low peptide coverage (32).

While bottom-up proteomics is still the preferred method for characterizing proteomes due to its coverage, robustness across diverse protein physiochemical properties, sensitivity, and quantitative capabilities --there remain challenges. Moving forward we will need new or repurposed methods, tools, and datasets to better interpret peptide level measurements. Datasets with known differences in peptide measurements will be crucial for validating any new approaches that are proposed to deal with peptide level differences. Additionally, improved data visualization tools are necessary to better distinguish changes inclusive of conserved domains, known PTMs, and structural features within a protein coding gene in the context of a global proteome. Finally, a compiled reference or “atlas” of experimentally-observed proteoforms presents a major opportunity for future algorithm development, which the Human Proteoform Atlas recently framed (36). As the technology has advanced, so too has our ability to obtain robust measurements across many samples without lots of missing data. We now need to move towards understanding why these peptide measurements may be different instead of simply forcing our data into a format in which it may not be best served and instead into a format in which it fits.

## Acknowledgements

This work was supported in part by National Institute of Health grants U19AG065156 (MJM, TJM, DLP), RF1AG053959 (MJM, TJM, DLP), P41GM103533 (MJM, DLP), F31AG069420 (DLP), P41GM108569 (NLK), UH3CA246635 (NLK), R35GM126914 (LMS), by Swedish Research Council grant 2017-04030 (LK), and U01–1CA184783 (BMW, LB) at PNNL, a multiprogram national laboratory operated by Battelle for the U.S. Department of Energy under contract DE-AC06-76RL01830. Some of the tissues used to produce the proteomic data in Figure 4 were provided by the Adult Changes in Thought study (R13AG057087).

